# The Ketogenic Diet Fails to Mitigate Seizures and Neuroinflammatory Responses in a Mouse Model of Virus-Induced Epilepsy

**DOI:** 10.64898/2026.05.19.726056

**Authors:** Casey H. Meili, Kaitlyn Allen, Daniel J. Doty, Sofia Del Fiol, Ana Beatriz DePaula-Silva

**Affiliations:** Department of Pharmacology and Toxicology, University of Utah, Salt Lake City, UT, 84112, USA

## Abstract

**Objective:** The ketogenic diet (KD) is a high-fat, low-carbohydrate intervention widely used to treat drug-resistant epilepsy, thought to reduce seizures through a combination of metabolic, neuronal, and microbiota-dependent mechanisms. Additionally, recent studies suggest that the anticonvulsant effects of KD require the gut microbiota, with taxa such as *Akkermansia* and *Parabacteroides* contributing to seizure protection by modulating host neurotransmitter balance and neural excitability. While KD has been shown to be effective in reducing seizure burden across different epilepsies, its antiseizure effect on infection-driven seizures, which are often driven by acute neuroinflammation, has not been evaluated. Here, we evaluated the effects of KD on seizure burden, neuroimmune responses, and gut microbiota composition in the Theiler’s murine encephalomyelitis virus (TMEV) model of virus-induced epilepsy.

**Methods:** Mice were maintained on either a KD or a normal diet prior to intracerebral TMEV infection. Seizures were induced by handling and scored twice daily from day 3 to 7 post-infection. Neuroimmune responses were assessed by flow cytometry, and fecal microbial composition was analyzed using 16S rRNA gene sequencing.

**Results:** Despite achieving ketosis, KD did not reduce seizure incidence, seizure burden, or seizure severity during acute TMEV infection. KD also did not significantly alter overall immune cell infiltration into the central nervous system, indicating limited effects on global neuroinflammation. However, KD significantly reshaped the gut microbiota, reducing alpha diversity (richness, Shannon diversity, and evenness) and strongly altering community structure with clear separation between diet groups, including enrichment of taxa such as *Akkermansia, Acetatifactor, Dorea*, and *Flintibacter*, and depletion of fiber-associated taxa including *Bifidobacterium* and *Roseburia*. However, these microbial shifts were insufficient to mitigate inflammation-driven seizures.

**Significance:** These results demonstrate that KD’s anticonvulsant efficacy is highly context-dependent, and that KD-driven changes in microbiota- and metabolite-mediated mechanisms may be ineffective against infection-associated epilepsy, suggesting that inflammation-driven seizures require distinct therapeutic approaches.

**Key points:**

- The ketogenic diet (KD) does not reduce acute seizure incidence and severity during TMEV infection despite achieving ketosis
- KD does not induce neuroinflammatory changes associated with seizure outcomes
- KD strongly reshapes gut microbiota, reducing diversity and altering community structure.
- Microbiota changes are insufficient to protect against inflammation-driven seizures
- KD anticonvulsant effects are context-dependent and ineffective in infection-driven epilepsy

## Introduction

Epilepsy is a chronic and debilitating neurological disorder affecting over 70 million people worldwide^1^. Despite the availability of numerous antiseizure medications (ASMs), approximately one-third of patients remain refractory to treatment, highlighting the need for alternative therapeutic strategies^2^. Neuroinflammation is increasingly recognized as a key contributor to seizure susceptibility and epileptogenesis. Central nervous system (CNS) infections, drive neuroinflammation and are a significant risk factor for temporal lobe epilepsy (TLE), the most common form of acquired epilepsy ^3^. A broad range of pathogens, including bacteria, fungi, parasites, and over 100 viruses, can infect the brain and cause encephalitis^4,5^. Patients who survive viral encephalitis are approximately 16 times more likely to develop epilepsy than the general population^6^, and those who experience acute symptomatic seizures during infection are 22 times more likely to develop epilepsy^7^, underscoring the need to understand how viral infection triggers seizures and identify strategies to prevent post-infectious epilepsy.

The Theiler’s murine encephalomyelitis virus (TMEV) model of virus-induced epilepsy recapitulates key features of TLE. Following intracerebral infection of TMEV, C57BL/6J mice develop acute symptomatic seizures 3-8 days post-infection (dpi), associated with robust neuroinflammatory responses, including the activation of microglia and infiltration of peripheral immune cells, particularly monocytes/macrophages^8,9^. This is followed by a latent phase where the virus is cleared, and seizures are no longer observed. Following the latent phase, a subset of mice that experience acute seizures later develop spontaneous recurrent seizures, modeling the progression to post-infectious epilepsy^10^. This chronic phase is characterized by hippocampal sclerosis and atrophy, gliosis, and peripheral immune cell infiltration into the brain, features that mirror those seen in human TLE^11,12^. These features make the TMEV model highly relevant for studying the mechanisms of infection-associated epilepsy and for testing candidate interventions aimed at reducing seizure burden and neuroinflammation.

The ketogenic diet (KD) is a non-pharmacological high-fat, low-carbohydrate, moderate protein dietary therapy that has demonstrated broad efficacy against multiple seizure types, often succeeding in patients who fail to respond to antiseizure medications when used as an adjunct to existing ASMs^13^. By heavily restricting carbohydrates, the KD shifts metabolism toward fatty acid oxidation, inducing a metabolic state known as ketosis. During ketosis, high levels of acetyl-CoA drive hepatic ketogenesis and the production of ketone bodies, primarily β-hydroxybutyrate and acetoacetate^14^. These ketone bodies circulate to the brain, where they serve as alternative energy substrates^15^. The KD is hypothesized to exert its anti-seizure effects through multiple, complementary mechanisms. First ketosis increases the ATP/ADP ratio, upregulates genes involved in mitochondrial metabolism, and promotes mitochondrial biogenesis in hippocampal neurons, thereby increasing neuronal energy reserves and resilience to metabolic stress^16–18^. Second, KD modulates neuronal excitability by increasing GABAergic signaling and reducing glutamatergic transmission, with ketone bodies capable of directly dampening excitatory neurotransmission in some animal models^19,20^. Third, KD has been shown to exert anti-inflammatory effects via β-hydroxybutyrate, which inhibits the NLRP3 inflammasome and reduces the production of pro-inflammatory cytokines, such as IL-1β, IL-6, and TNF-α^21^. These anti-inflammatory effects are partly mediated by microglia and infiltrating macrophages, which adopt a less proinflammatory reactive state under ketotic conditions^22,23^.

More recently, the gut microbiota has emerged as a critical mediator of KD efficacy. KD induces profound shifts in microbial community composition, and these changes have been linked to seizure protection in both preclinical models and clinical studies^24–26^. Notably, Olson *et al*. demonstrated that depletion of the gut microbiota abolishes the anticonvulsant effects of KD in mouse models of both electrically and chemically induced seizures (6 Hz and pentylenetetrazole (PTZ)), while reconstitution with KD-associated microbial communities restores seizure protection^26^. Specific taxa, including *Akkermansia muciniphila* and *Parabacteroides* spp., were shown to be necessary for this protective effect and to modulate host metabolic and neurotransmitter pathways, including increased hippocampal γ-aminobutyric acid (GABA) levels and an altered glutamate-GABA balance^26^. These findings suggest that KD may exert anticonvulsant effects, in part, through microbiota-dependent modulation of neural excitability and metabolism.

However, whether these mechanisms are relevant in the context of infection-driven seizures, and whether KD can reduce acute seizure burden or modulate the neuroinflammatory response, has not been studied. Filling this gap will clarify KD’s therapeutic potential for infection-associated epilepsy and its effects on seizure susceptibility during neuroinflammation. Therefore, in this study, we aimed to determine whether ketogenic diet treatment reduces acute seizure burden and neuroinflammation in the TMEV model of viral encephalitis. In addition, we assessed the impact of KD on the gut microbiota to determine whether diet-induced microbial changes are associated with altered seizure outcomes in this model. We hypothesized that KD would attenuate seizure frequency and severity, potentially through combined effects on host metabolism, immune responses, and microbiota composition.

## Results

### The ketogenic diet fails to reduce seizure incidence or severity

To determine the effect of KD in modulating seizures induced by viral infection, mice were fed either a ketogenic diet (KD) or a normal diet (ND) for 38 days prior to intracerebral TMEV infection and monitored for seizures from 3 to 7 days dpi (Fig 1A). Seizure incidence was comparable between groups, with 13/20 ND mice and 12/21 KD mice exhibiting at least one seizure during the acute seizure phase (p = 0.7513, Fisher’s exact test) (Fig 1B). Daily cumulative seizure burden also did not differ between KD and ND mice (p > 0.05, two-way ANOVA with Sidak’s multiple comparisons test) (Fig 1C). Furthermore, there was no significant difference in the number of seizures per mouse (Mann-Whitney test, p = 0.4025) (Fig 1D). Importantly, a shorter-duration KD paradigm (10 days on KD diet prior to infection; 17 days total at sacrifice) yielded similar results, with no differences in seizure incidence, cumulative seizure burden, or number of seizures per mouse (Fig S1). To confirm ketosis, plasma ketone body levels were quantified using a colorimetric β-hydroxybutyrate (β-HB) assay at the day of infection (n=5 per group) and at 7 dpi (n=12 per group) after 45 days on diet. At both time points, KD-fed mice demonstrated significantly greater β-HB levels compared to ND controls (Fig S2), confirming sustained ketosis throughout the experimental period. Together, these findings indicate that KD treatment did not alter acute seizure susceptibility or severity in the TMEV-infected mice.

**Figure 1.**
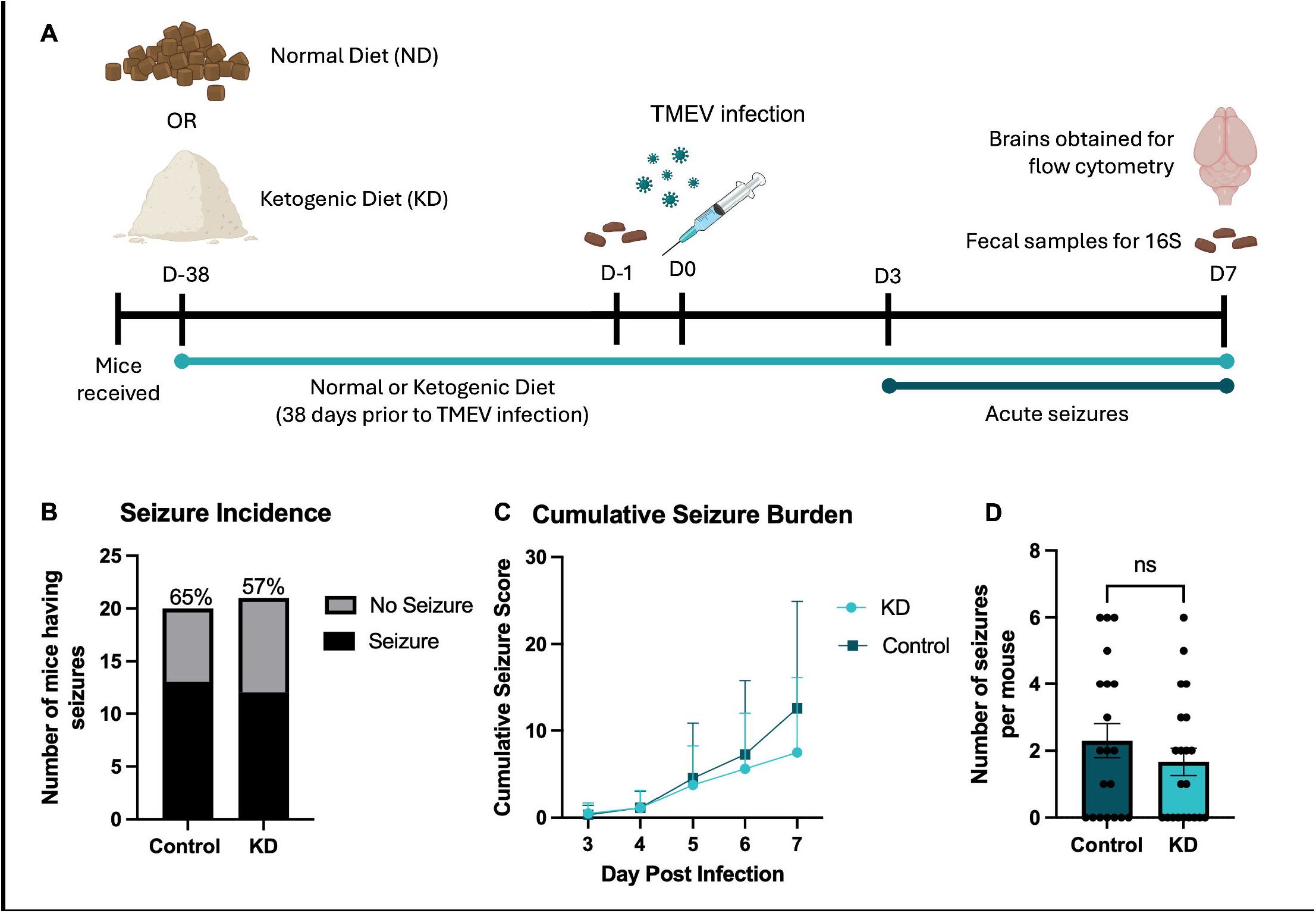
KD treatment fails to reduce seizure incidence or severity in TMEV-infected mice. (A) Experimental design schematic. Mice were maintained on either a ketogenic diet (KD) or normal diet (ND) control for 38 days prior to intracerebral TMEV infection. Seizures were monitored twice daily from 3-7 days post-infection. Fecal samples were collected at baseline (day -1 relative to infection) and at 7 dpi for 16S rRNA gene sequencing. At 7 dpi, mice were euthanized for flow cytometric analysis of brain immune cell populations. (B) Total number and percentage of mice with observed seizures (Control: 13/20; KD: 12/21); Fisher’s exact test, p = 0.7513. (C) Daily cumulative seizure burden; two-way repeated-measures ANOVA with Sidak’s multiple-comparisons test; all p values > 0.05. (D) Number of observed seizures per mouse; data are shown as mean ± SEM, with each point representing an individual mouse; Mann-Whitney test, p = 0.4025.

### Ketogenic diet alters lymphocyte composition without broadly affecting myeloid activation during TMEV infection

Flow cytometry analysis was performed on cells isolated from half-brains at 7 dpi of TMEV-infected mice fed KD or ND to evaluate cell content and neuroinflammatory response. Cell populations were identified using standard gating strategies (Fig S3). There were no significant differences between KD- and ND-fed mice in the total number and percentage of infiltrating cells (CD45^high^) infiltrating macrophages, lymphocytes, CD8^+^ T cells, and resident microglia (Welch’s t-test or Mann-Whitney U test, p > 0.05 for all comparisons) (Fig 2A-E). However, KD-fed mice exhibited a significant reduction in NK1.1^+^ cells as a percentage of total lymphocytes compared to ND controls (mean ± SEM, KD = 18.78 ± 1.416%; ND = 25.84 ± 1.478%; Welch’s t-test p = 0.0039) (Fig 2F), indicating modulation of natural killer cell populations. In addition, KD-fed mice showed a significant increase in CD4^+^ T cells as a percentage of total lymphocytes compared to ND controls (mean ± SEM, KD = 15.93 ± 1.548%; ND = 11.62 ± 1.106%; Mann-Whitney test p = 0.0148) (Fig 2G). Assessment of activation markers on microglia and brain-infiltrating monocytes/macrophages revealed no significant differences in MHC-II, TREM1, or TREM2 expression (p > 0.05 for all comparisons) (Fig 2H-K). Additionally, a trend toward reduced Ly6C expression on macrophages was observed in KD-fed mice, although this did not reach statistical significance (Welch’s t-test, p = 0.0607) (Fig 2L). In the shorter-duration KD cohort, we similarly observed a significant reduction in NK1.1+ cells and a decrease in macrophage Ly6C expression, with no differences in macrophage, microglia, lymphocyte or CD4^+^ T cell percentages and no changes in MHC-II or TREM2 expression (Fig S4). These findings suggest that KD modulates lymphocyte composition without broadly altering myeloid cell activation in the TMEV model.

**Figure 2.**
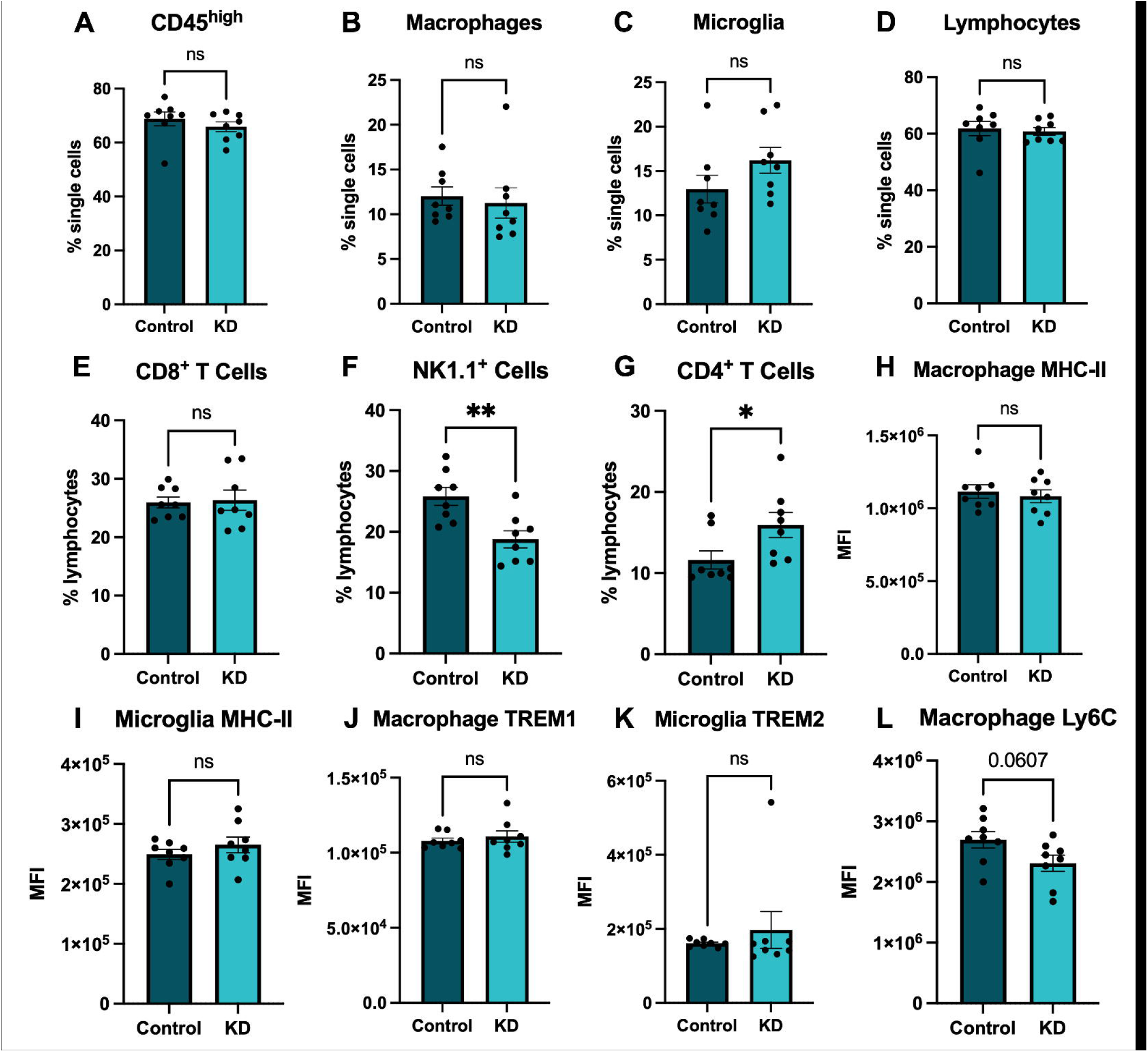
Ketogenic diet alters lymphocyte proportions in the mouse brain during TMEV infection. Flow cytometric analysis of brain immune cells at 7 dpi. Cells were gated on live, singlet populations prior to downstream analysis. Microglia and macrophages were identified based on CD45 and CD11b expression, with microglia defined as CD45^low/int^ CD11b^+^ and macrophages defined as CD45^high^ CD11b^+^. Lymphocytes were gated as CD45^+^ CD11b^−^ cells. (A) Total CD45^high^ cells as a percentage of single cells. (B) Macrophages as a percentage of single cells. (C) Microglia as a percentage of single cells. (D) Lymphocytes as a percentage of single cells. (E) CD8^+^ T cells as a percentage of lymphocytes. (F) NK1.1^+^ cells as a percentage of lymphocytes. (G) CD4^+^ T cells as a percentage of lymphocytes. (H) Mean fluorescence intensity (MFI) of MHC-II on macrophages. (I) MFI of MHC-II on microglia. (J) MFI of TREM1 on macrophages. (K) MFI of TREM2 on microglia. (L) MFI of Ly6C on macrophages. Each point represents an individual mouse (n = 8 per group; 4 seizure, 4 non-seizure), and data are shown as mean ± SEM. Statistical significance was determined using Welch’s t-test (C-F, H-J, L) or Mann-Whitney test (A-B, G, K) for each panel: (A) p = 0.2345, (B) p = 0.3282, (C) p = 0.1504, (D) p = 0.7364, (E) p = 0.8447, (F) p = 0.0039, (G) p = 0.0148, (H) p = 0.6151, (I) p = 0.3287, (J) p = 0.5161, (K) p = 0.3823, (L) p = 0.0607.

### Ketogenic diet reshapes gut microbiota composition and diversity

Previous studies have shown that KD significantly modulates gut microbiota composition; therefore, to determine the effect of KD on gut microbial communities during acute seizure development in TMEV-infected mice,16S rRNA sequencing was performed on fecal samples collected from control ND and KD-fed mice at baseline (day -1 relative to TMEV infection, following 38 days on diet) and at 7 dpi to assess the impact of KD on gut microbial communities (Fig. 3). Sequencing depth was sufficient to capture community diversity across all samples (average coverage = 99.5%; Supplementary Table 1). Across both groups, the gut microbiota was dominated by members of the Lachnospiraceae family; however, distinct compositional differences were observed between diets. KD-fed mice exhibited higher relative abundances of *Akkermansia, Flintibacter, Lawsonibacter*, and *Acetatifactor*, whereas ND mice were enriched in *Paramuribaculum* and *Bifidobacterium* (Fig. 3A). Alpha diversity analyses revealed significant differences between diets at both time points. KD-fed mice displayed reduced richness (Sobs) compared to ND-fed controls at day -1 (p = <0.0001) and 7 dpi (p = <0.0001) (Fig. 3B). Similarly, KD mice exhibited significantly lower Pielou’s evenness at day -1 (p = 0.0074) and 7 dpi (p = 0.0411) (Fig. 3C), as well as reduced Shannon diversity at both time points (day -1 p = 0.0003; 7 dpi p = 0.0025) (Fig. 3D), indicating an overall reduction in microbial diversity associated with KD. Beta diversity analysis using Bray-Curtis dissimilarity demonstrated clear separation between KD and ND control groups at both day -1 and 7dpi, indicating distinct microbial community structures (Fig. 3E). These differences were statistically significant (PERMANOVA R^2^ = 0.572, p = 0.001), consistent with a strong effect of diet on microbiota composition.

**Figure 3.**
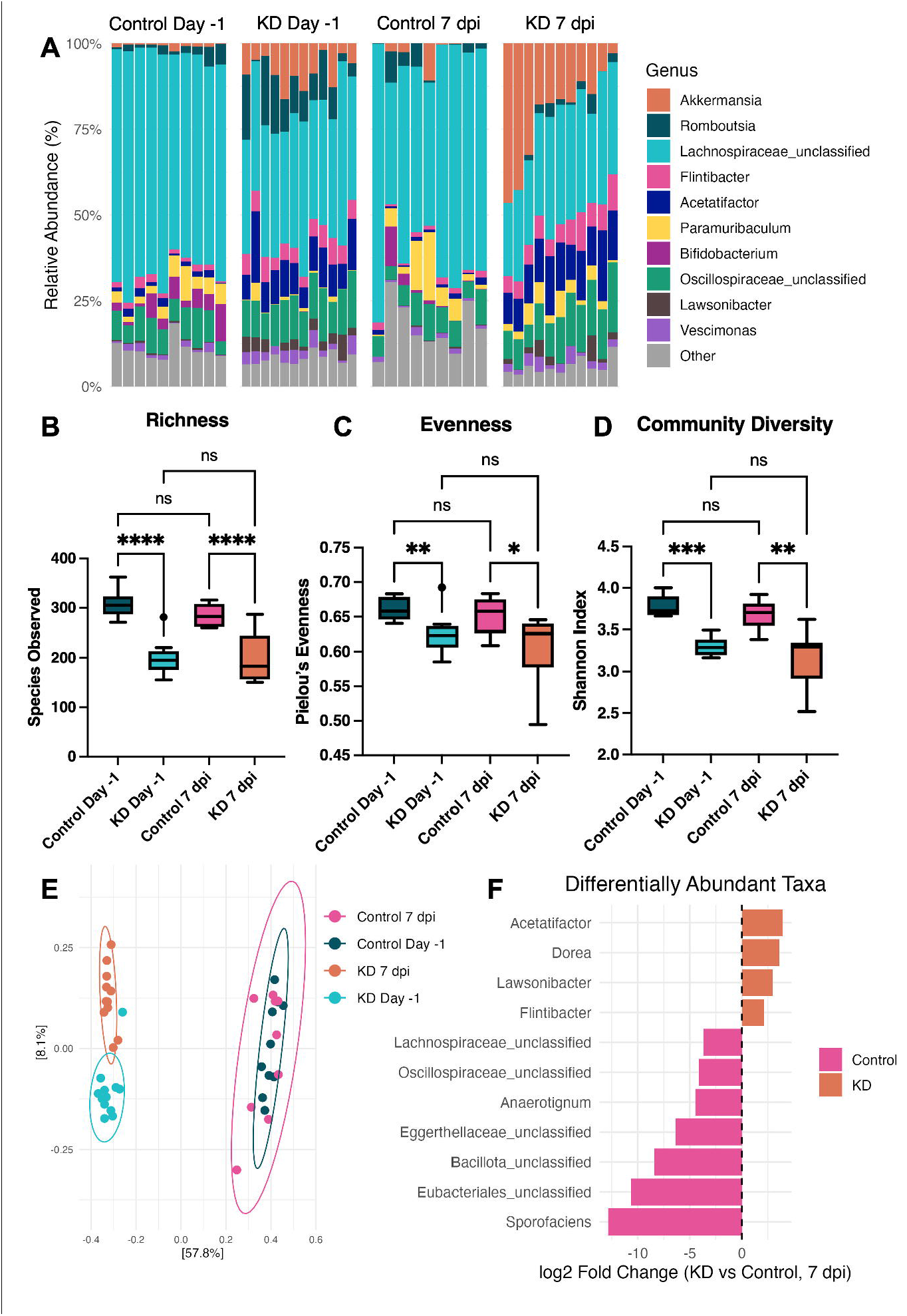
KD modulates gut microbial diversity and community structure. Fecal samples were collected from ND-fed control and KD-fed mice at baseline (day -1, after 38 days on diet) and 7 days post infection for 16S rRNA gene sequencing. All analyses were performed on TMEV-infected mice only. (A) Relative abundance of the 10 most abundant bacterial taxa at baseline and 7 dpi, displayed as stacked bar plots at the genus level where available. (B) Observed species richness (Sobs), (C) Pielou’s evenness, and (D) Shannon diversity index at day -1 and 7 dpi. Data are shown as Tukey box plots (center line, median; box, interquartile range; whiskers, 1.5x IQR), Comparisons were performed using ordinary one-way ANOVA (sobs) or Kruskal-Wallis (Pielou’s evenness and Shannon) tests with post hoc multiple-comparisons correction. (E) Principal coordinates analysis (PCoA) based on Bray-Curtis dissimilarity illustrating microbial community structure across diet and infection groups. Each point represents an individual mouse, with 95% confidence ellipses shown for each group. Group differences were assessed using PERMANOVA (KD vs. ND control at both time points, PERMANOVA; R^2^ = 0.57202, p = 0.001), and homogeneity of dispersion was evaluated using betadisper. (F) Differential abundance analysis at 7 days post-infection (7 dpi) between KD and ND control groups was performed using DESeq2 and visualized as shrinkage-adjusted log_2_ fold change (KD vs ND). The 10 taxa with the largest absolute log_2_ fold changes are shown. Taxa were aggregated at the genus level based on taxonomic classification; when genus-level assignment was not possible, the lowest available taxonomic level (e.g., family-level unclassified taxa) was retained. Positive values indicate enrichment in the KD group, and negative values indicate enrichment in the ND control group.

Differential abundance analysis at 7 dpi using DESeq2 identified multiple taxa significantly enriched between groups (Fig. 3F, Supplementary Table 2). KD-fed mice were enriched in taxa including *Acetatifactor* (log_2_FC = 2.97, adjusted p = 1.85 x 10^-27^), *Flintibacter* (log_2_FC = 1.66, adjusted p = 2.34 x 10^-12^), *Dorea* (log_2_FC = 3.31, adjusted p = 1.89 x 10^-6^), and *Lawsonibacter* (log_2_FC = 2.61, adjusted p = 3.56 x 10^-4^), and *Akkermansia* (log_2_FC = 2.90, adjusted p = 9.6 x 10^-3^), a taxon previously implicated in ketogenic diet–associated seizure protection. In contrast, ND-fed mice were enriched in unclassified Eggerthellaceae (log_2_FC = -7.08, adjusted p = 7.36 x 10^-38^), *Sporofaciens* (log_2_FC = -8.70, adjusted p = 2.21 x 10^-21^), unclassified Lachnospiraceae (log_2_FC = - 1.66, adjusted p = 7.03 x 10^-14^), and *Clostridium sensu stricto* (log_2_FC =-7.65, adjusted p = 1.08 x 10^-9^), as well as enrichment of fiber-associated and saccharolytic taxa including *Roseburia* (log_2_FC = - 13.44, adjusted p = 7.10 × 10^-38^), *Bifidobacterium* (log_2_FC = -11.58, adjusted p = 1.25 × 10^-20^), and *Muribaculaceae* (log_2_FC = -11.83, adjusted p = 1.15 × 10^-14^). Together, these findings demonstrate that KD induces significant shifts in specific bacterial taxa during TMEV infection.

## Discussion

The ketogenic diet is a well-established therapy for drug-resistant epilepsy; however, its efficacy across diverse seizure models remains uncharacterized. Here we demonstrate that KD does not reduce seizure incidence or severity in the TMEV model of infection-induced epilepsy. Despite achieving ketosis, KD-fed mice exhibited seizure incidence and seizure burdens comparable to ND-fed animals. Notably, this lack of efficacy was consistent across both short- and long-term KD paradigms, indicating that insufficient duration of dietary intervention does not explain the absence of seizure protection. These findings indicate that the anticonvulsant effects of KD do not extend to this model of acute, virally driven seizures. The lack of efficacy observed here contrasts with the well-documented benefits of KD in genetic and chemoconvulsant models of epilepsy^27–30^. TMEV-induced seizures are driven by acute neuroinflammation, which may override or bypass the neuronal and metabolic pathways through which KD is thought to exert its anticonvulsant effects. Together, these findings support the idea that KD efficacy is highly context-dependent and may be limited in settings of infection-driven seizures.

Additionally, because acute seizures in the TMEV model are associated with robust neuroinflammatory responses, including infiltration of peripheral immune cells into the CNS and activation of macrophages and microglia^31–33^, we evaluated the impact of KD on the neuroimmune response during acute infection. While KD did not alter the overall magnitude of immune cell infiltration into the CNS, it did shift lymphocyte composition with a reduction in NK1.1^+^ cells and an increase in CD4^+^ T cells. This reduction in NK1.1+ cells is consistent with findings in cancer models, where KD has been shown to reduce NK cell abundance and impair function^34^. These findings suggest that KD can modulate specific immune cell subsets without broadly suppressing neuroinflammation. Natural killer cells play roles in response to viral infections and in inflammation^35^, though they do not drive seizure development in this model, as NK1.1-depleted mice show no difference in seizure incidence^36,37^. The lack of an effect on seizures despite reduced NK1.1^+^ cells confirms that modulation of this population alone is insufficient to alter disease severity. Although KD increased the proportion of CD4+ T cells, prior studies have demonstrated that CD4+ T cells do not drive seizure development in this model, indicating that this population is not a determinant of disease outcome in this context^37,38^. In addition, KD-fed mice exhibited a trend toward reduced infiltration of Ly6C^hi^ macrophages, suggesting a potential shift toward a less inflammatory phenotype, although this did not reach statistical significance. Together, these findings indicate that while KD may subtly influence immune cell phenotype, these effects are insufficient to alter seizure outcomes in this model.

In addition to its effects on immune responses, KD induced substantial changes to the gut microbiota. KD-fed mice exhibited reduced richness, evenness, and overall diversity relative to ND-fed controls, alongside clear shifts in community composition and differential enrichment of specific taxa. These findings are consistent with prior studies, in both rodent models and patients, demonstrating that KD imposes strong ecological constraints on the gut microbiome, favoring taxa adapted to low-carbohydrate, high-fat environments^39–41^. Consistent with this, KD selectively enriched taxa such as *Acetatifactor, Flintibacter, Dorea*, and *Lawsonibacter*, while ND mice were enriched in taxa including *Sporofaciens, Clostridium sensu stricto*, and members of the Lachnospiraceae and Eggerthellaceae families. Many of the KD-enriched taxa identified here fall within Clostridial lineages implicated in fermentation, including production of short-chain fatty acids and other bioactive metabolites that can influence immune function and gut-brain signaling^42^. However, the extent to which these taxa produce functionally relevant metabolites under ketogenic conditions remains unclear, particularly given reduced dietary carbohydrate availability. Despite significant microbiome changes, KD did not confer protection against seizures in the TMEV model, indicating that ketogenic diet-induced alterations in microbial community structure alone are insufficient to modify acute seizure outcomes in this context.

Several studies have suggested that the anticonvulsant effects of KD are mediated through the gut microbiota. In particular, Olson *et al*. demonstrated that KD protection against both electrically induced (6 Hz) and chemoconvulsant (PTZ) seizures requires an intact microbiota, and that enrichment of *Akkermansia* and *Parabacteroides* drives this effect^26^. These taxa were shown to be associated with increased hippocampal GABA levels and altered hippocampal glutamate:GABA ratios, and transplantation of KD-associated microbiota was sufficient to confer seizure protection in recipient mice^26^. In our study, KD robustly reshaped the gut microbiota and enriched several taxa previously linked to KD-associated metabolic outputs, yet this did not translate into reduced seizure susceptibility in TMEV-infected mice. Notably, while *Akkermansia* was enriched, we did not observe enrichment of *Parabacteroides*, a key taxon implicated in microbiota-mediated seizure protection, suggesting that its absence may contribute to the lack of efficacy. However, KD instead enriched alternative taxa, including *Acetatifactor, Dorea*, and *Lawsonibacter*, which may share overlapping functional capacities, such as fermentation and production of bioactive metabolites that influence host metabolism and gut-brain signaling^43–45^. This raises the possibility that functionally similar, but taxonomically distinct, microbial communities were established in our model, which may partially recapitulate KD-associated metabolic function without being sufficient to confer seizure protection. This contrast suggests that while microbiota shifts may be necessary for KD-mediated anticonvulsant effects, they are not sufficient in the context of acute, inflammation-driven seizures. It is possible that microbial metabolites sufficient to protect against chemoconvulsant-induced seizures do not overcome the strong neuroinflammatory and viral mechanisms driving TMEV seizures. These findings emphasize that microbiota-dependent mechanisms of KD are likely context-specific, and that these shifts alone may not guarantee seizure protection in infection-associated epilepsy.

Importantly, limitations of this study should be considered. First, our analyses were focused on the acute phase of TMEV infection and therefore do not address whether KD may influence the development or severity of spontaneous recurrent seizures during the chronic epileptic phase. Given that epileptogenesis involves distinct mechanisms from acute seizure generation, future longitudinal studies are needed to determine whether KD treatment can modify long-term outcomes. Second, while 16S rRNA sequencing identified significant diet-induced changes in microbial composition, this approach does not resolve functional details like microbial metabolite production, which are likely critical mediators of gut-brain interactions. Future studies incorporating metabolomics or shotgun metagenomics will be necessary to determine whether KD-associated microbial communities influence disease outcomes in a context-dependent manner. The lack of efficacy observed here suggests that metabolic and microbiota-mediated mechanisms alone may be insufficient to overcome the strong neuroinflammatory drivers of TMEV-induced seizures, highlighting the need for combinatorial or etiology-specific therapeutic strategies targeting both inflammation and neuronal excitability.

In summary, KD induces robust metabolic and microbiome changes and selectively alters immune cell composition yet fails to reduce seizure incidence or severity in a viral infection driven model. These findings highlight important limitations in the generalizability of KD as an anticonvulsant therapy and emphasize the need to consider underlying disease mechanisms when evaluating metabolic interventions. More broadly, this work suggests that effective treatment of infection-associated epilepsy may require etiology-specific approaches.

## Methods

### Mouse husbandry

All mouse experiments were performed in compliance with the guidelines approved by the University of Utah Institutional Animal Care and Use Committee. 6-week-old male C57BL/6J mice were obtained from Jackson Laboratory (Bar Harbor, ME, USA) and housed together (5 per cage) in a facility providing a 12-hour light/dark cycle. Food and water were provided *ad libitum*. Only male mice were used as sex-dependent differences have not been reported in this model^46^.

### Ketogenic diet

Mice cages were randomized and received either the control normal diet (ND) or ketogenic diet (KD) *ad libitum*, beginning 38 days prior to TMEV infection. Teklad 2920X (Inotiv, Madison, WI, USA) served as the control diet. The ketogenic diet was custom formulated and irradiated by Teklad (Inotiv, Madison, WI, USA) and contained 44% Crisco, 15% Cocoa Butter, and 8.5% corn oil. The ketogenic diet was stored at 4°C. Ketosis was evaluated over the course of the experiment using ketone urine test strips (Keto-Check Inc, Napa, CA, USA) and plasma β-Hydroxybutyrate on day -3 and 7 days post infection

### β-Hydroxybutyrate (ketone body) colorimetric assay

Blood samples were obtained with heparinized needles by cardiac puncture at the time of sacrifice (day -3 and 7 dpi) and centrifuged at 12,000 rpm for 30 minutes at 4°C. Plasma was removed from the pellet. For the mice sacrificed at 7 dpi, the plasma was stored at -80°C until the time of assay. For the mice sacrificed prior to infection, assay was run immediately after using non frozen/thawed plasma. Samples were analyzed using the β-Hydroxybutyrate (Ketone Body) Colorimetric Assay Kit (cat. #700190) from Cayman Chemical (Ann Arbor, MI, USA) according to the manufacturer’s instructions. Absorbance at 450 nm was measured. Samples were run in triplicate and averaged. β-Hydroxybutyrate concentrations were calculated using the standard curve and kit instructions.

### TMEV infection and seizure scoring

Mice were briefly anesthetized using isoflurane and injected with 20μl of either sterile phosphate-buffered saline (PBS) (n=5 KD, n=5 ND) or 20μl PBS containing 4.08 x 10^5^ plaque-forming units (PFUs) DA-TMEV (n=25 KD, n=25 ND) intracortically in the right hemisphere using an insulin syringe fitted with a William’s collar to limit needle penetration to 2 mm. Seizures were induced by handling mice twice on days 3-7 post-infection, with handling sessions separated by at least 6 hours. Seizures were scored using the modified Racine scale as follows: stage 1 – mouth and facial movements, stage 2 – head nodding, stage 3 – forelimb clonus, stage 4 – forelimb clonus and rearing, stage 5 – forelimb clonus, rearing, and falling, and stage 6 – jumping, running, repeated falling, and severe clonus^31^. The percentage of mice with seizures was calculated as follows: (number of mice with seizures / total number of mice infected) × 100. The cumulative seizure burden was calculated by summing all seizure scores for each mouse.

### Brain tissue dissociation and flow cytometry

Following euthanasia with isoflurane, mice were perfused with PBS, and brains were harvested. Half brains (contralateral to injection site) were finely chopped with a razor blade, added to 2ml of digestion solution containing RPMI and 1mg/ml of collagenase D (Sigma-Aldrich, St. Louis, MO, USA), and incubated for 20 minutes at 37°C. Cells in collagenase were homogenized by manually pipetting against a petri dish. The petri dish was washed with RPMI, and the homogenate was then passed through a 100 μm filter. Following centrifugation at 500g and 4°C for 7 minutes, cells were resuspended in 37% Percoll (Sigma-Aldrich, St. Louis, MO, USA) and spun again for 20 minutes at 700g. Myelin and supernatant were then aspirated. Cells were resuspended in a buffer containing 1X PBS and 3% fetal bovine serum (FBS) and then filtered again through a 40 μm filter. Cell suspensions were counted by trypan blue exclusion, and approximately 500,000-viable cells were plated per well in 96-well V-bottom plates for flow cytometry staining.

For flow cytometry analysis, Fc receptors were first blocked using mouse FcR Blocking Reagent (Miltenyi Biotec, San Diego, CA, USA), as indicated by the manufacturer. The cells were then incubated with their respective cell-surface antibodies in 100 µl of PBS + 3% FBS for 15 minutes at 4°C. The following fluorochrome-labeled antibodies were used: CD11b (PerCP/Cy5.5, clone M1/70, BioLegend, cat #101228, 1:100), CD45 (V500, clone 30-F11, BD Biosciences, cat #561487, 1:100), Ly-6C (APC/Fire750, clone HK1.4, BioLegend, cat #128046, 1:100), MHC-II (Alexa Fluor 488, clone M5/114.15.2, BioLegend, cat #107616, 1:100), NK1.1 (BUV661, clone PK136, BD Biosciences, cat #741477, 1:100), CD4 (BV785, clone RM4-5, BioLegend, cat #100552, 1:100), CD8a (Alexa Fluor 700, clone 53-6.7 BioLegend, cat #100730, 1:100), TREM1 (Pe, R&D Systems, cat #FAB1187P 1:10), and TREM2 (APC, R&D Systems, cat #FAB17291A, 1:10). 5µl of Super Bright Complete Staining Buffer (Invitrogen, Carlsbad, CA, USA) was added to all samples to prevent non-specific fluorochrome interactions. After staining, cells were washed and resuspended in PBS with 3% FBS for acquisition. Samples were acquired on a NovoCyte Penteon Flow Cytometer (Agilent Technologies, Santa Clara, CA, USA) with the instrument set to record either 70 μL of sample or 50,000 P2 events (single cells), whichever occurred first. Flow cytometry data were first gated on live cells (P1) and subsequently on singlets (P2). From the singlet population, microglia were defined as CD45^low/int^ CD11b^+^ cells, while infiltrating macrophages were identified as CD45^high^ CD11b^+^ cells. Lymphocytes were defined as CD45^high^ CD11b^−^ cells and further characterized into subsets, including CD4^+^ T cells, CD8^+^ T cells, and NK1.1^+^ cells. Downstream analyses were performed using NovoExpress Software. Representative flow plots showing the gating strategy are shown in Fig S2. Fluorescence minus one (FMO) controls were used to establish gating boundaries for each marker (CD45, CD11b, CD4, CD8, NK1.1, TREM1, TREM2, MHC-II, Ly6C). Compensation for spectral overlap was performed using single-stained compensation beads (BioLegend, cat #424602) according to the manufacturer’s instructions. All mean fluorescence intensity (MFI) values are reported as geometric mean fluorescence intensity.

### 16S sequencing and amplicon analysis

DNA was extracted from fecal samples using the ZymoBIOMICS DNA Miniprep Kit (Zymo Research, Irvine, CA, USA) according to the manufacturer’s instructions, with a 40-minute manual homogenization using a vortex adaptor. The V3-V4 region of the 16S rRNA gene was amplified using the Quick-16S Plus NGS Library Prep Kit (V3-V4, UDI) (Zymo Research, Irvine, CA, USA). Each sample was assigned a unique dual index (UDI) barcode. Following PCR amplification, products were pooled by equal volume and purified prior to sequencing. All library preparation and purification steps were done according to the manufacturer’s protocol. The final pooled library was quantified, assessed for quality, and sequenced on an Illumina MiSeq (2 x 300 bp paired-end reads with a PhiX spike-in) at the University of Utah’s High-Throughput Genomics Shared Resource (Salt Lake City, UT, USA).

16S sequencing data were processed using mothur v.1.48.2^47^, using a modified MiSeq SOP (accessed March 2026)^48^. Paired-end reads were assembled into contigs, and low-quality sequences were removed based on length (400-460 bp), ambiguous bases, and homopolymer stretches (>8 bp). Duplicate sequences were merged, and the remaining sequences were aligned to a customized SILVA v138.2 reference database trimmed to the V3-V4 region. Sequences that failed to align within the expected region were discarded. After filtering, sequences were denoised by pre-clustering (allowing 2 bp differences) and screened for chimeras using VSEARCH. Taxonomic classification was performed using the RDP reference database (trainset19, release 2023), and non-bacterial lineages (chloroplasts, mitochondria, archaea, eukaryotes, and unclassified sequences) were removed. Sequences were clustered into operational taxonomic units (OTUs) at 97% similarity. Consensus taxonomy was assigned to OTUs, and a subsampled OTU table was generated to account for uneven sequencing depth.

### Alpha and beta diversity

Alpha diversity was assessed in mothur. To account for uneven sequencing depth across samples, the OTU table was subsampled to the second smallest group size (18,670 sequences per sample). The sample below this cutoff was discarded from all subsequent analyses. Diversity metrics (observed species richness (sobs), Shannon diversity, and pielou’s evenness) were then calculated using the summary.single command.

Beta diversity analyses were performed with R (v4.4.1) using the phyloseq package. OTU, taxonomy, and metadata tables generated in mothur were imported into a phyloseq object. Bray-Curtis dissimilarities were calculated, and ordination was performed using principal coordinates analysis (PCoA). Significance of differences between groups was tested using permutational multivariate analysis of variance (PERMANOVA; adonis2 function in vegan), and homogeneity of group dispersions was assessed using betadisper^49^. Ordination plots were visualized with ggplot2, with 95% confidence ellipses overlaid to illustrate group clustering.

OTU abundance, taxonomic assignments, and sample metadata were used to construct a phyloseq object in R (v4.4.1) using the phyloseq package^50^. Community compositional differences between groups were assessed using Bray-Curtis dissimilarity. Principal coordinates analysis (PCoA) was performed using the ordinate function in phyloseq. Ordination plots were generated with samples colored by experimental group, and 95% confidence ellipses were added to visualize group clustering. Visualization was performed using ggplot2. To test for differences in microbial community structure between experimental groups, PERMANOVA was performed using the adonis2 function from the vegan package with Bray-Curtis distance matrices. Homogeneity of group dispersions was assessed using the betadisper function, followed by significance testing using both ANOVA and permutation tests to determine if group differences were driven by differences in dispersion.

### Differential abundance analysis

Differential abundance testing was conducted using DESeq2^51^ within the phyloseq framework. First, samples were subset to the 7 dpi. Prior to differential abundance testing, OTUs were agglomerated to the genus level using taxonomic classification. Low-abundance taxa were filtered by removing features with total counts ≤ 10 across all samples. Count data were modeled using a negative binomial generalized linear model implemented in DESeq2 with diet as the explanatory variable. The reference level was set to control 7 dpi, enabling interpretation of positive log2 fold changes as enrichment in the KD group. Statistical inference (p-values and FDR-adjusted p-values) and reported effect sizes in the Results section were derived directly from DESeq2 outputs.

To improve visualization and reduce the influence of extreme log2 fold change estimates in low-count taxa, shrinkage of log2 fold changes was additionally performed using apeglm^52^. These shrinkage-adjusted values were used exclusively for figure generation (Fig. 3F) to provide more conservative and interpretable effect size estimates for plotting. Differential abundance results were merged with taxonomic classifications from the phyloseq taxonomy table. Taxa were consistently defined at the genus level; sequences not classified to genus were retained as higher-level placeholders (e.g., “Family_unclassified”). Taxa were considered statistically significant at a false discovery rate (FDR)-adjusted p-value < 0.05. For visualization purposes, the top 10 differentially abundant genera were selected based on the magnitude of shrinkage-adjusted log2 fold change and displayed as a bar plot indicating directionality and relative effect size between groups.

### Statistics

All statistical analyses, except those used for beta diversity and microbiome differential abundance, were performed using GraphPad Prism 11 (GraphPad Software, San Diego, CA, USA). A p-value of < 0.05 was used to determine significance for all statistical tests. Seizure incidence was compared using Fisher’s exact test. Total number of observed seizures per mouse was analyzed using the Mann-Whitney U test, and daily cumulative seizure burden was analyzed using two-way repeated measures ANOVA with Sidak’s multiple comparisons test. Flow cytometry data were analyzed using Welch’s t-tests or Mann-Whitney tests, depending on data normality.

Alpha diversity was assessed in mothur after rarefaction of the OTU table to 18,670 sequences per sample, and one sample below this threshold was excluded from downstream analysis. Diversity metrics (observed species richness, Shannon diversity index, and Pielou’s evenness) were compared between groups using ordinary one-way ANOVA (for normally distributed metrics) or Kruskal-Wallis tests (for non-parametric metrics), followed by appropriate post hoc corrections. Beta diversity and differential abundance analyses were conducted using R v4.4.1. Beta diversity was evaluated using Bray-Curtis dissimilarity. Group differences in microbial community composition were assessed using permutational multivariate analysis of variance (PERMANOVA; adonis2 function in vegan). Homogeneity of multivariate dispersion was tested using betadisper followed by ANOVA and permutation testing to ensure that observed differences were not driven by dispersion effects.

For differential abundance analysis, genus-level taxonomic counts from 7 days post-infection samples were analyzed using DESeq2 within the phyloseq framework. A negative binomial generalized linear model was fitted with diet as the main explanatory variable, with control samples at 7 dpi set as the reference condition. Low-abundance taxa (total counts ≤ 5 across all samples) were removed prior to modeling. Log2 fold changes were shrinkage-adjusted using the apeglm method to improve effect size estimation and reduce variance inflation for low-count taxa^52^. These shrinkage-adjusted values were used exclusively for data visualization. Taxa were considered statistically significant at an FDR-adjusted p-value < 0.05. For visualization, the top differentially abundant taxa were selected based on the largest absolute shrinkage-adjusted log2 fold change.

## Data availability statement

The sequencing data generated in this study have been deposited in the NCBI Sequence Read Archive (SRA) under BioProject accession number PRJNA1459435 and are available under SRA accession numbers SRR38321467-SRR38321521.

## Code availability

All code used for 16S sequencing processing and downstream analyses is available on GitHub at: https://github.com/caseymeili/Ketogenic-Diet-TMEV. This repository includes the mothur pipeline used for sequence processing and OTU table generation, as well as R scripts used for phyloseq-based community analyses, including Bray-Curtis dissimilarity calculations, PCoA ordination, PERMANOVA (adonis2), dispersion testing (betadisper), and DESeq2-based differential abundance analysis.

## Author Contributions

Casey Meili: Formal Analysis, Investigation, Data Curation, Writing – Original Draft, Writing – Review & Editing, Visualization

Kaitlyn Allen: Formal Analysis, Investigation, Visualization Daniel Doty: Investigation Sofia Del Fiol: Investigation

Ana Beatriz DePaula Silva: Conceptualization, Writing – Review & Editing, Supervision, Funding Acquisition

