## Supplementary Figures 1-4 for "The Ketogenic Diet Fails to Mitigate Seizures and Neuroinflammatory Responses in a Mouse Model of Virus-Induced Epilepsy"

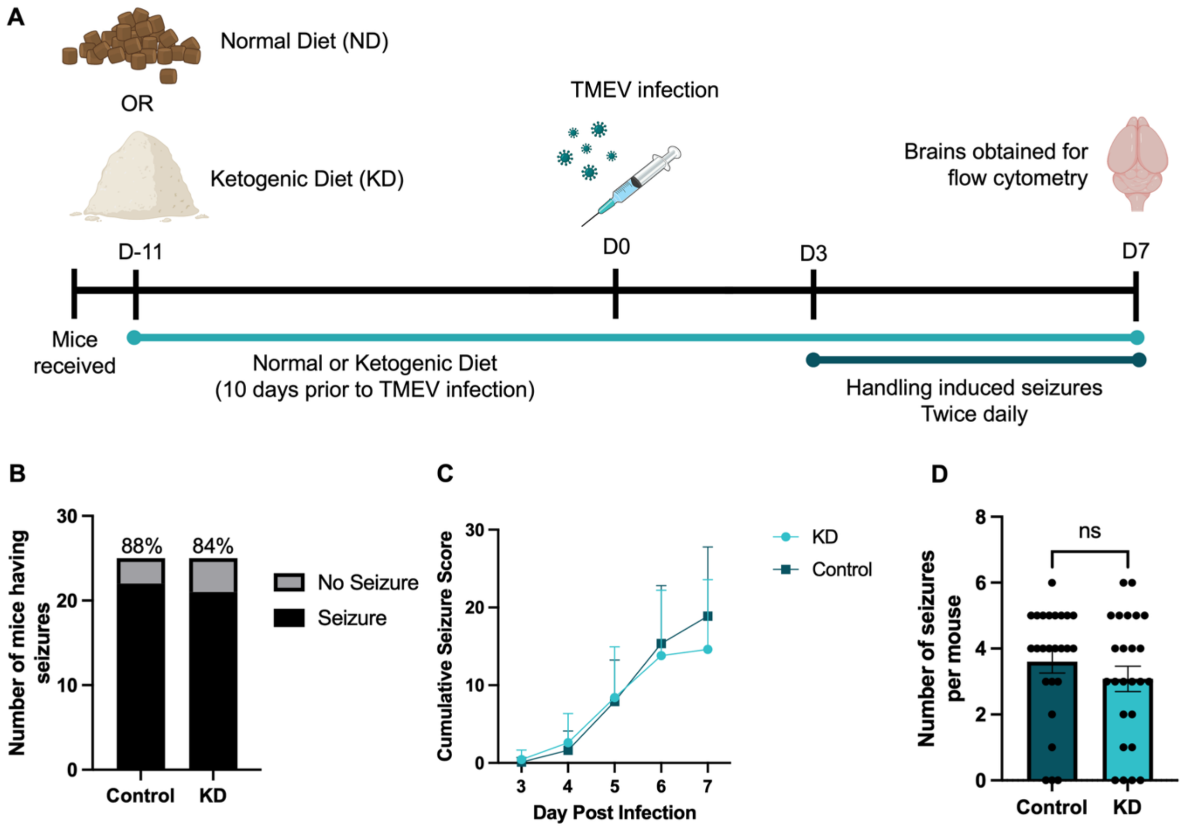


**Supplementary Figure 1 – Short-term KD treatment fails to reduce seizure incidence or severity in TMEV-infected mice.** (A) Experimental design schematic. Mice were maintained on either a ketogenic diet (KD) or normal diet (ND) control for 10 days prior to intracerebral TMEV infection. Seizures were monitored twice daily from 3-7 days post-infection. At 7 dpi, mice were euthanized for flow cytometric analysis of brain immune cell populations. (B) Total number and percentage of mice with observed seizures (Control: 22/25; KD: 21/25); Fisher's exact test, p = > 0.999. (C) Daily cumulative seizure burden; two-way repeated-measures ANOVA with Sidak's multiple-comparisons test; all p values > 0.05. (D) Number of observed seizures per mouse; data are shown as mean ± SEM, with each point representing an individual mouse; Mann-Whitney test, p = 0.2849.

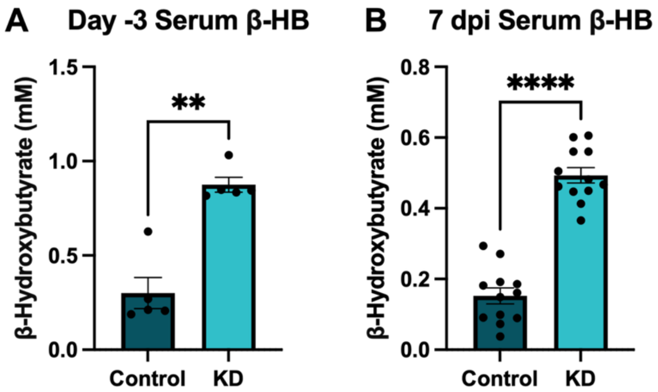


**Supplementary Figure 2 – Ketogenic diet induces ketosis in mice.** Plasma β-hydroxybutyrate (BHB) levels (mM) were measured in mice maintained on control or ketogenic diet (KD). (A) Baseline BHB levels at day -3 prior to infection, after 38 days on KD (no freeze–thaw; n = 5 per group). (B) BHB levels at 7 days post-infection from plasma samples subjected to freeze-thaw (n = 12 per group). Data are presented as mean ± SEM. Statistical significance was determined using (A) Mann-Whitney Test p = 0.0079, or (B) Welch's t-test p = <0.0001.

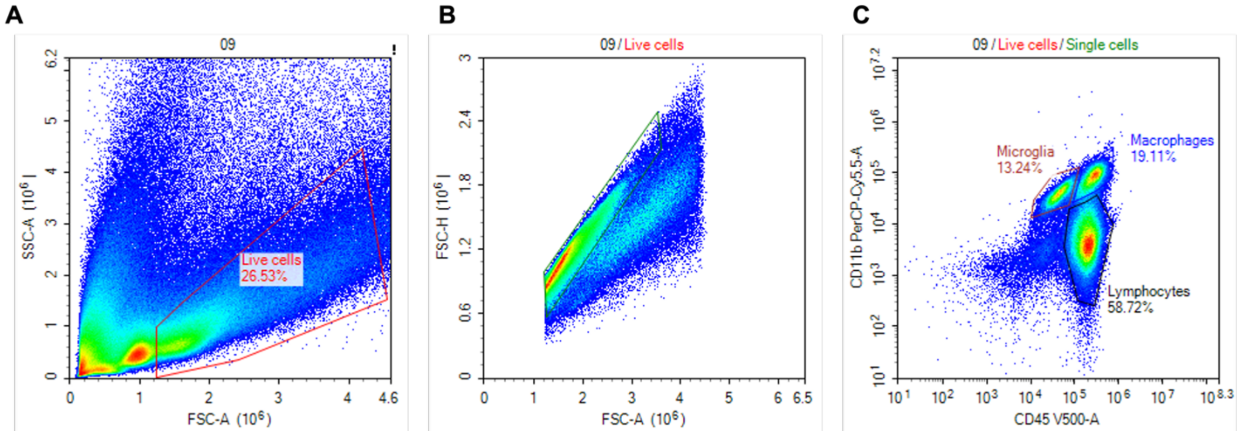


**Supplementary Figure 3 – Flow cytometry gating strategy for identification of brain immune cell populations.** Representative gating strategy used for analysis of brain single-cell suspensions. (A) Initial gating on side scatter area (SSC-A) and forward scatter area (FSC-A) was used to exclude debris and dead cells. (B) Singlets were gated based on FSC-A vs. FSC-height (FSC-H) to exclude doublets. (C) CD45 versus CD11b expression was used to distinguish major immune cell populations, with microglia defined as $CD45^{low/int}$ $CD11b^{+}$, macrophages defined as $CD45^{high}$ $CD11b^{+}$, and lymphocytes defined as $CD45^{+}$ $CD11b^{-}$. Fluorescence minus one (FMO) controls were used to establish gating boundaries for each marker (CD4, CD8, NK1.1, TREM1, TREM2, MHC-II, Ly6C)

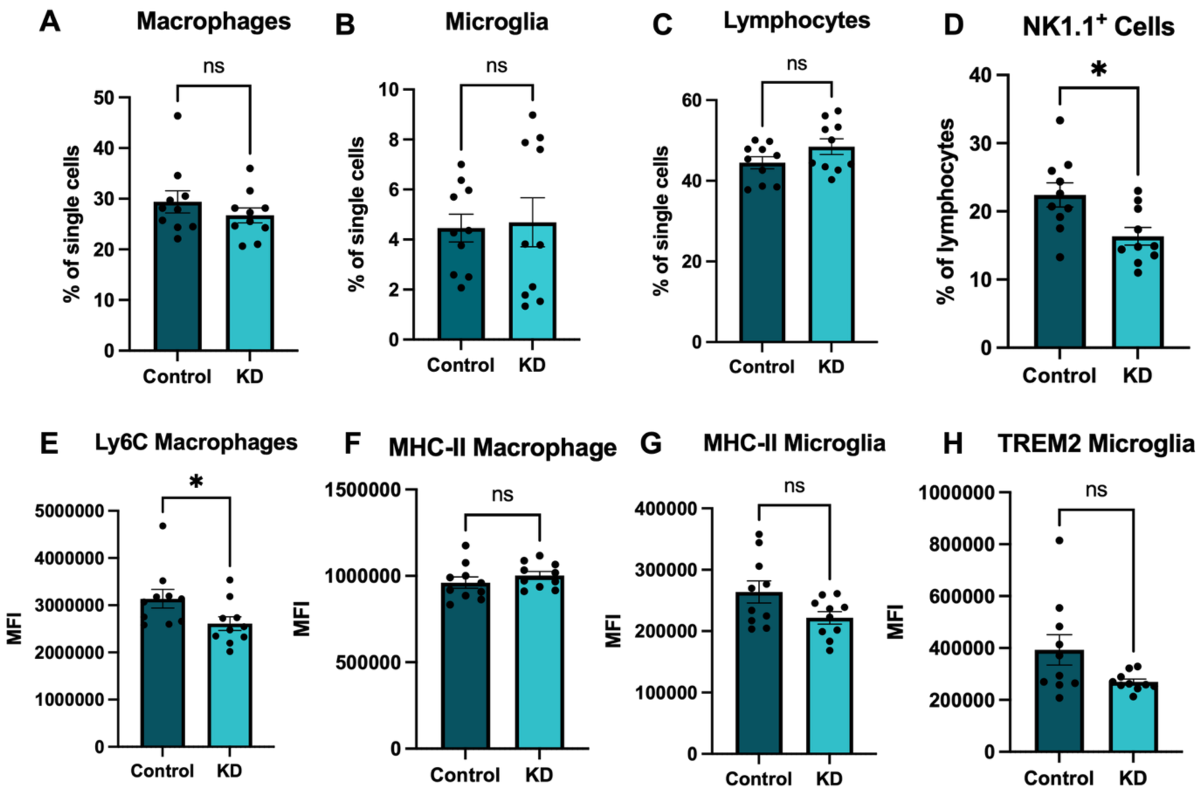


**Supplementary Figure 4 – Short-term ketogenic diet feeding does not alter brain immune cell composition or activation during TMEV infection.** Mice were maintained on the ketogenic diet for 10 days prior to infection with TMEV and remained on diet until tissue collection at 7 dpi (17 total days on diet). Flow cytometric analysis of brain immune cells was performed at 7 dpi. Cells were gated on live, singlet populations prior to downstream analysis, as described in Figure 2, Methods, and Supplementary Figure 2. (A) Macrophages as a percentage of single cells; Mann-Whitney test; $p = 0.4353$. (B) Microglia as a percentage of single cells; Mann-Whitney test; $p = 0.9705$. (C) Lymphocytes as a percentage of single cells; Welch's t-test; $p = 0.1233$. (D) NK1.1$^+$ cells as a percentage of lymphocytes; Welch's t-test; $p = 0.0131$. (E) Mean fluorescence intensity (MFI) of Ly6C on macrophages; Mann-Whitney test; $p = 0.0355$. (F) MFI of MHC-II on macrophages; Welch's t-test; $p = 0.3225$. (G) MFI of MHC-II on microglia; Welch's t-test; $p = 0.0598$. (H) MFI of TREM2 on microglia; Welch's t-test; $p = 0.0649$. Each point represents an individual mouse ($n = 10$ per group), and data are shown as mean ± SEM.

**Supplementary Table 1 – Sample metadata, sequencing depth, and alpha diversity metrics.** Table includes all 16S samples analyzed, along with associated metadata, total number of sequences per sample, sequencing coverage, and alpha diversity indices, including observed richness (Sobs), Pielou's evenness (Shannon evenness), and Shannon diversity.

**Supplementary Table 2 – Differentially abundant taxa.** Differential abundance analysis of fecal bacterial taxa between KD and normal diet-fed TMEV-infected mice at 7 days post-infection determined using DESeq2. Taxa with adjusted p values (Benjamini-Hochberg corrected) < 0.05 are shown. Positive $log_2$ fold change values indicate enrichment in KD-fed mice, while negative values indicate enrichment in ND-fed mice. Taxonomic assignments are shown from domain to genus level where available.
